# Epitope-Based Peptide Vaccine against Bombali Ebolavirus Viral Protein 40: An Immunoinformatics Combined with Molecular Docking Studies

**DOI:** 10.1101/2020.12.28.424637

**Authors:** Mujahed I. Mustafa, Shaza W. Shantier, Miyssa I. Abdelmageed, Abdelrafie M. Makhawi

**Affiliations:** Department of Biotechnology, University of Bahri, Khartoum, Sudan; Department of Pharmaceutical Chemistry, Faculty of Pharmacy, University of Khartoum, Khartoum, Sudan; Faculty of Pharmacy, University of Khartoum, Khartoum, Sudan

**Keywords:** Bombali Ebolavirus, matrix protein, Viral Protein (VP40), vaccine design, Immunoinformatics

## Abstract

**Background:** Bombali Ebolavirus is RNA viruses belong to the Filoviridae family. They are causing lethal hemorrhagic fever with high mortality rate. Despite having available molecular knowledge of this virus, no approved vaccine or antiviral drugs have been developed yet for the eradication of Bombali Ebolavirus infections in humans.

**Objective:** the present study described a multi epitope-based peptide vaccine against Bombali Ebolavirus matrix protein VP40, using several immunoinformatics tools.

**Materials and Methods:** The six strains of Ebolavirus were retrieved from NCBI and Uniprot databases and submitted to VaxiJen to identify the most antigenic protein among all. Then PSIPRED, SOPMA, QMEAN, and PROCHECK tools were used to check the protein quality. T-cell prediction, population coverage, and molecular docking analysis were achieved to select peptides containing multiple Bombali VP40 epitopes showing interaction with multiple HLA molecules for expected immune response across the world.

**Result:** Bombali Ebola (YP_009513276.1) was found to be the most antigenic protein among all. Which it has been used in all required analysis. For T cell three epitopes showed high affinity to MHC class I (YSFDSTTAA, VQLPQYFTF, and MVNVISGPK) and high population coverage against Africa and the world. Furthermore in MHC class II, six promising epitopes that associated with most common MHC class II alleles.

**Conclusion:** The above result conclude that, these peptides capable of provoking T-cell response and being interacted with a wide range of HLA molecules have a strong potential to be a vaccine against Bombali Ebolavirus.

## 1 Introduction

Ebola viruses are RNA viruses belong to the *Filoviridae* family [1, 2]. They are causing lethal hemorrhagic fever with high mortality rate [3-5]. The main source of the ebolavirus is unknown, but several mammal groups are the source from which it is transferred to the human species [6]. In addition to Africa, it has been reported in the United Kingdom, and Italy, and secondary cases (Health-care workers) have occurred in the United States and Spain [7, 8].

It characterized by an immediate onset of flu-like illness, followed a preliminary incubation period of 2–21 days. After this preliminary period of infection, the symptoms became noticeable which include chest pain, cough, nausea, vomiting and mucosal hemorrhage [9].

So far, six different strains of EBOV have been identified: Sudan (SEBOV), Ivory Coast, Zaire (ZEBOV), Reston, Bundibugyo and the most recently identified strain Bombali virus. Among them, ZEBOV is considered as the most frequently occurring infection with the highest number of deaths. Ebola viruses infection is considered to be the most deadly one having mortality rate as high as 90% in human cases only [10]. Due to this high mortality rates, lack of treatment options and vaccination make Ebola virus a biothreat pathogen [11].

The Ebola virus matrix protein VP40 (viral protein 40 kDa) is the most abundantly expressed filoviral protein which serves as the primary matrix protein and coordinates virion assembly at the plasma membrane through interactions with both viral and cellular components. [12]. The matrix protein VP40 with their novel adjuvant have already been demonstrated to be safe when administered intramuscularly or subcutaneously, and therefore, they are closer to clinical trials than adjuvants whose safety profiles are unknown [13]. With the available knowledge, several attempts have been made to produce affective vaccine against Ebola viruses due to its deadly nature.

Vaccine design methods based on T-cell epitope can be characterized as the recognition of immune-dominant epitopes of the virus and synthesizing it to be utilized as vaccines to induce effective immune response. These assessments enhance the possibility of an ideal vaccine candidate. Computer-based prediction tools reduce the number of validation experiments and time for epitope prediction. The method of epitope-based vaccine design has been performed against several life-threating diseases [14-17]. Although, most of epitope-based vaccines are developed based on B cell epitopes, yet the potentiality of T cell epitope-based vaccine is also promising because CD8+ T cell can induce a more effective immune response of the host cell towards the infected T cells [18].

Development of an effective and safe Ebola virus vaccine has been hindered by a lack of knowledge. In this study, we attempted to identify major immunogenic epitopes on Ebola viral proteins and predict a vaccine. Simultaneously, we also achieved a genome wide search to recognize the most appropriate vaccine target site by using *in silico* tools. The study will enhance further upcoming laboratory-based approach to develop effective vaccine against Bombali Ebolavirus infection.

## 2. Material and methods

The immunoinformatics approaches used for vaccine design are illustrated in **Figure 1**.

**Figure 1.**
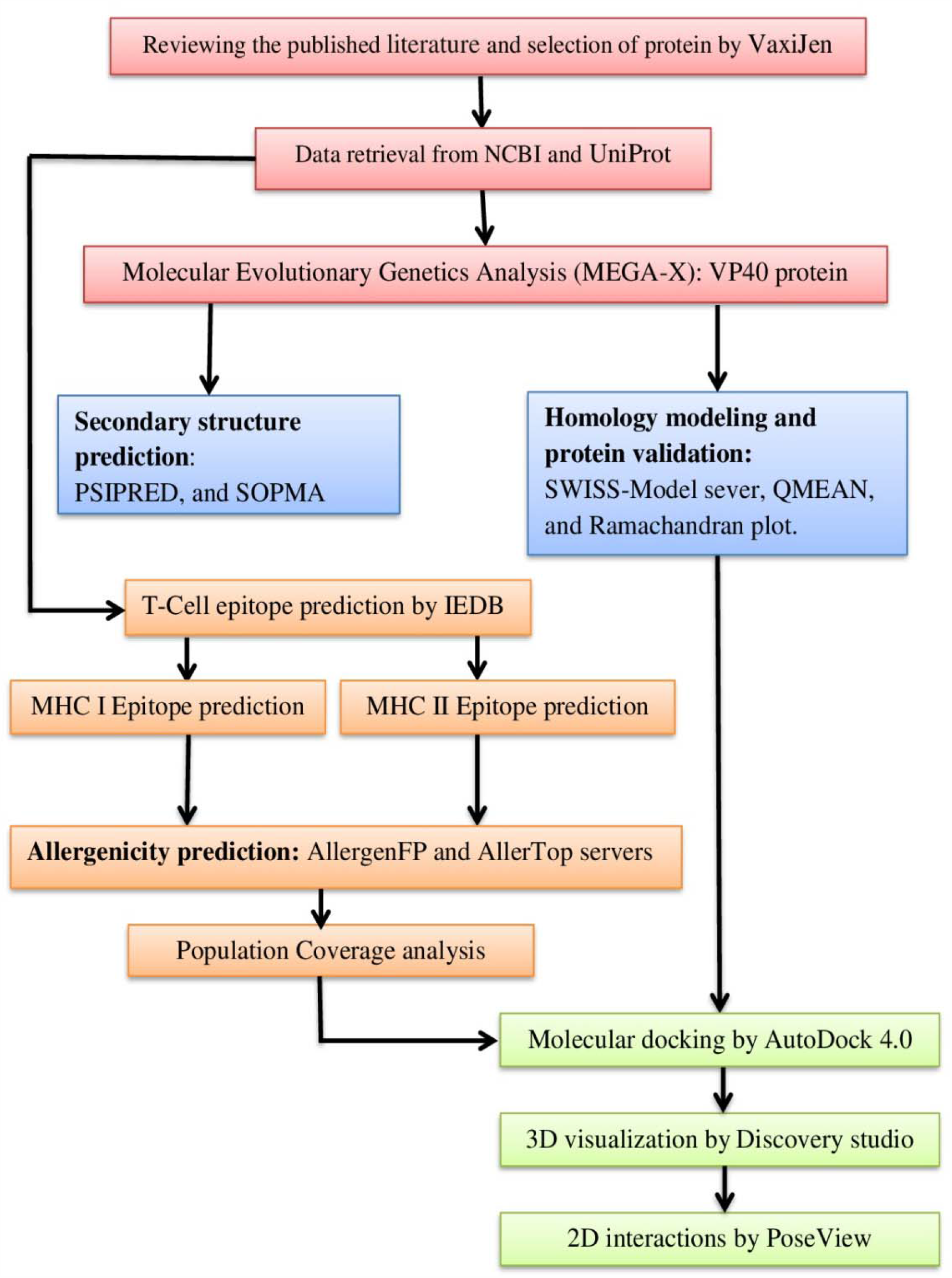
Illustrated the immunoinformatics approaches used for vaccine design against Bombali Ebolavirus VP40.

### 2.1. Sequence retrieval

The FASTA format of the reference sequence of Bundibugyo Ebola (NC_014373), Tai Forest Ebola (NC_014372), Zaire Ebola (NC_002549), Bombali Ebola (NC_039345), Sudan Ebola (NC_006432), and Reston Ebola (NC_004161), alongside the matrix protein (VP40) of Bombali Ebola (YP_009513276.1) were retrieved from the National Center of Biotechnology Information (NCBI) (https://www.ncbi.nlm.nih.gov/). The matrix protein (VP40) of Zaire ebolavirus (Q77DJ6), Tai Forest ebolavirus (B8XCN8), Reston ebolavirus (Q8JPX9), Bundibugyo ebolavirus (B8XCM9) and Sudan ebolavirus (Q5XX06) were retrieved from UniProt (https://www.uniprot.org/).

The 3D structures of MHC I allele HLA-A*02:01 and MHC II alleles HLA-DRB1*07:01, HLA-DPA1*02:01/DPB1*14:01 were retrieved from RCSB PDB (protein data bank) database. Swiss PDB viewer V.4.1.0 software [19] was used for structure optimization and energy minimization.

### 2.2. Antigenicity prediction

Antigenicity prediction of the matrix protein (VP40) of the six Ebola strains was performed by VaxiJen using threshold of 0.4 VaxiJen Server (http://www.ddg-pharmfac.net/vaxijen/VaxiJen/VaxiJen.html) allows the classification of antigens solely based on the physicochemical properties of proteins without referring to sequence alignment [20].

### 2.3. Conservation region analysis

Pairwise and multiple sequence alignment of the six strains of Ebola virus were performed using ClustalW. With gap opening penalty of 10.00 for both alignment, gap extension penalty of 0.10 for pairwise sequence alignment and 0.20 for multiple sequence alignment. Maximum likelihood phylogenetic tree was constructed with bootstrapping value of 300.

### 2.4. Phylogenetic analysis

Phylogenetic analysis reveals the evolutionary relationship between the different strains of the Ebola virus and trace back patterns of common ancestry between their lineages [21]. The analysis was performed using Molecular Evolutionary Genetics Analysis (MEGA-X) (version 10.1.18) (https://www.megasoftware.net). It is a software applied for sequence alignment, inferring phylogenetic trees, and testing evolutionary relationship. It can analyze DNA, RNA and Protein sequences [22].

### 2.5. Secondary structure prediction, homology modeling and protein validation

The secondary structures of the vaccine design were predicted using PSIPRED [23] by submitting the amino acids sequence as an input. PSIPRED is a secondary structure prediction online tool used to predict the transmembrane topology, transmembrane helix, fold and domain recognition. Its result was confirmed by another secondary structures online tool, SOPMA [24]. While the tertiary structure of Bombali Ebolavirus matrix protein was generated using SWISS-Model sever [19] because there is no structure available in Uniprot database [25]. Discovery studio 2020 [26] was used to visualize the most promising peptides for vaccine design. The confirmation of the predicted 3D structure was done using QMEAN [27], while the protein quality was tested using PROCHECK server by Ramachandran plot [28].

### 2.6. T-Cell epitope prediction tools

#### 2.6.1. Peptide Binding to MHC Class I Molecules

To predict the interaction with different MHC I alleles, the Major Histocompatibility Complex class I (MHC I) binding prediction tool on the IEDB http://tools.iedb.org/mhcI [29]. It services distinct approaches to measure the binding affinity of selected sequence to a definite MHC class I molecule. The half maximal inhibitory concentration (IC50) values of peptide binding to MHC class I molecule was calculated by artificial neural network (ANN) approach. All alleles having a binding affinity of IC50 that are equal or less than 500 nM were selected for further analysis.

#### 2.6.2. Peptide Binding to MHC Class II Molecules

The MHC II prediction tool provided by the IEDB was used to predict the peptide binding to MHC class II molecules, consisting of human allele references sets was used. The ANN prediction method was selected to recognize the binding affinity of MHC II grooves and MHC II binding core epitopes [30]. All epitopes that bind to many alleles at a score equal to or less than 1000, IC50 were chosen for additional investigation. The Human allele reference sets (HLA DR, DP, and DQ) were included in the prediction.

### 2.7. Allergenicity prediction

AllergenFP v.1.0 [31] and AllerTop v 2.0 servers [32] were used to evaluate the allergenicity of the predicted peptides.

### 2.8. Population coverage

The population coverage for each epitope was calculated by the IEDB population coverage tool at http://tools.iedb.org/tools/population/iedb_input [33]. All epitopes and their binding to MHC I and MHC II molecules were assessed against population covering the World and Africa.

### 2.9. Molecular docking analysis

In order to estimate the binding affinities between the proposed epitopes and molecular structure of T cells, in silico molecular docking was used. Sequences of proposed epitopes were selected from Ebola virus reference sequence using Chimera 1.10 and saved as PDB file. The obtained files were then optimized and energy minimized.

Molecular docking was then performed using AutoDock 4.0 software [34]. The active residue of the protein was selected, the results less than 1.0A□ in positional root-mean-square deviation (RMSD) were then considered ideal and clustered together for finding the favorable binding [35]. The highest binding energy (most negative) was considered as the ligand with maximum binding affinity. The 3D and 2D interactions of the resultant docking files with poses showing the lowest binding energies were visualized using DS Visualizer Client 2020 and the PoseView [36] at the ProteinPlus web portal [37], respectively.

## 3. Results

### 3.1. Conservation and antigenicity analysis

The multiple sequence alignment of the six strains of Ebola revealed eight conserved regions **(Table 1)**. Analysis of those conserved regions by VaxiJen revealed five sequences are antigenic as they met the criteria of default threshold ≥ 0.5 in VaxiJen **(Table 1)**.

**Table 1:** Predicted conserved regions with their antigenicity scores by VaxiJen

| Conserved peptide | start | End | VaxiJen score |
| --- | --- | --- | --- |
| LRPIADDNIDH | 51 | 61 | 0.2337 |
| KVLMKQIPIWLPLGV | 86 | 100 | 0.3173 |
| YSFDSTTAAIMLASYTITHFGK | 106 | 127 | 0.6932 |
| SNPLVRINRLG | 129 | 139 | 0.5018 |
| HPLRLLRIGNQAFLQEFVLPPVQLPQYFTFDLTALKLITQPLPAAT | 145 | 190 | 0.4827 |
| LRPGISFHPKLRPILLP | 203 | 219 | 1.5593 |
| KNIMGIEVPE | 256 | 265 | -0.1386 |
| IIPILLPKYIG | 284 | 294 | 0.6551 |

### 3.2. Phylogenetic analysis

The obtained multiple sequence alignment of the six Ebola virus strains was used to construct the phylogenetic tree shown in **Figure 2**.

**Figure 2:**
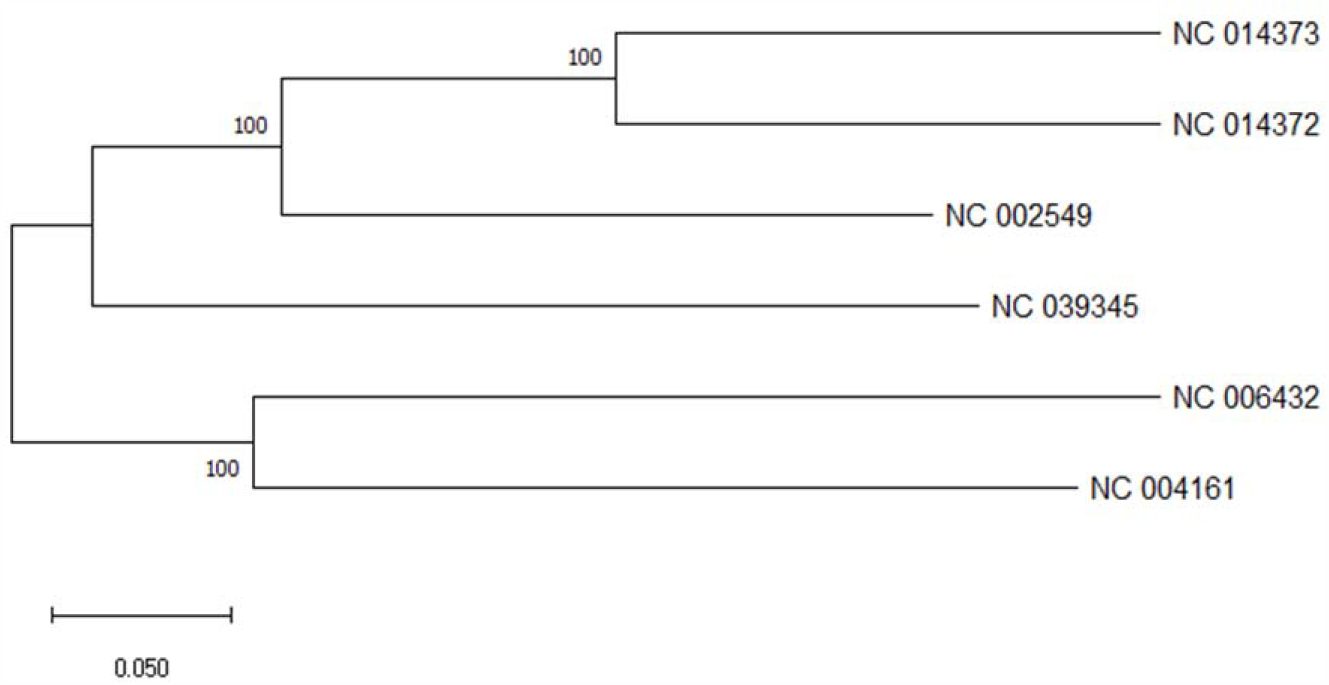
Illustrates the maximum likelihood phylogenetic tree, which constructed based on whole-genome sequence alignment of the six Ebola virus strains.

### 3.3. Secondary structure prediction

Out of the 326 amino acids analyzed by PSIPRED, 59 amino acids formed Alpha helix, 175 amino acids were involved in random coil, and 20 amino acids formed Beta turn; This result was confirmed by another secondary structures online tool, SOPMA **(Figures 3, 4)**.

**Figure 3:**
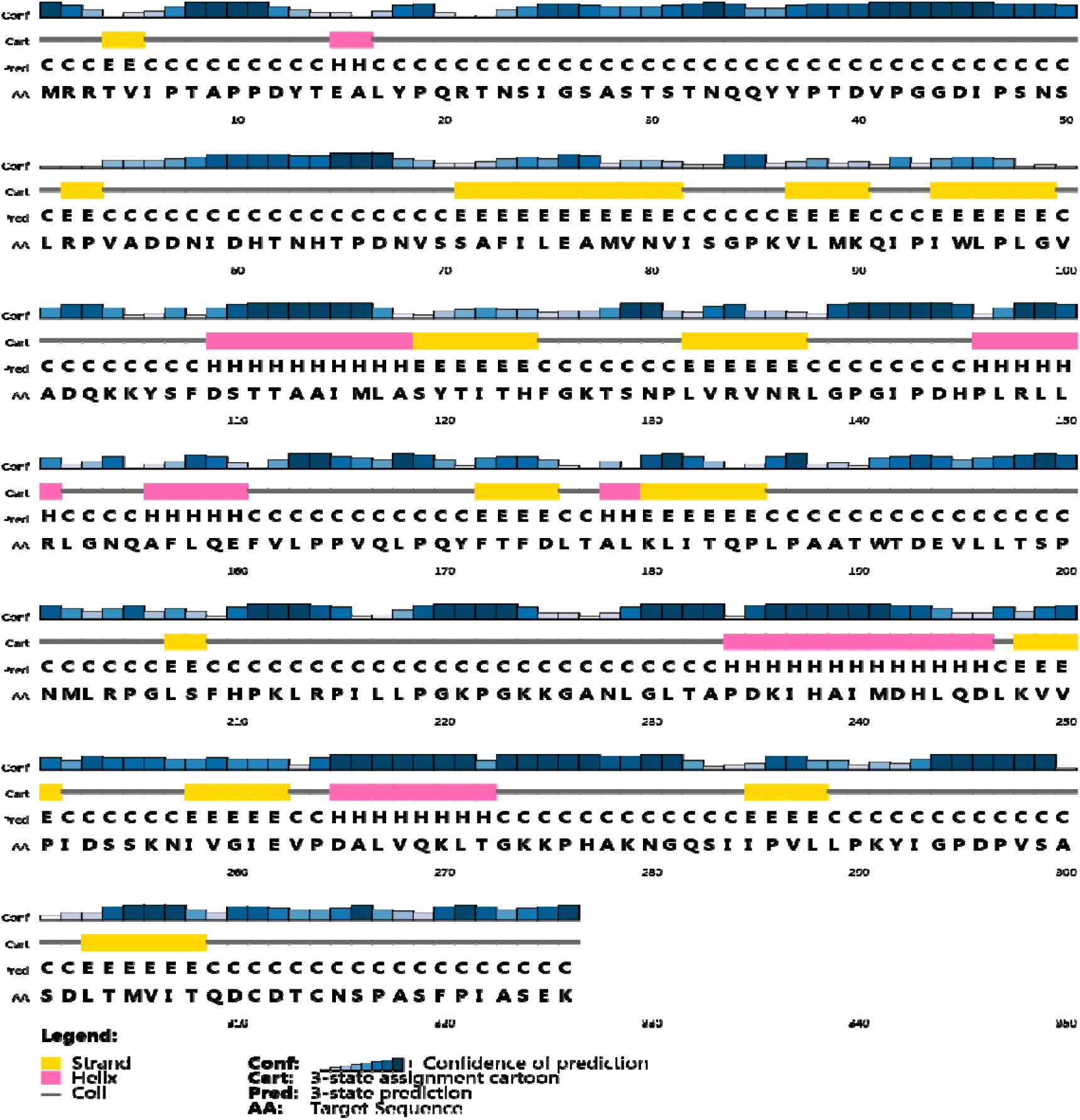
PSIPRED analysis of the secondary structure prediction of Bombali Ebolavirus matrix protein.

**Figure 4:**
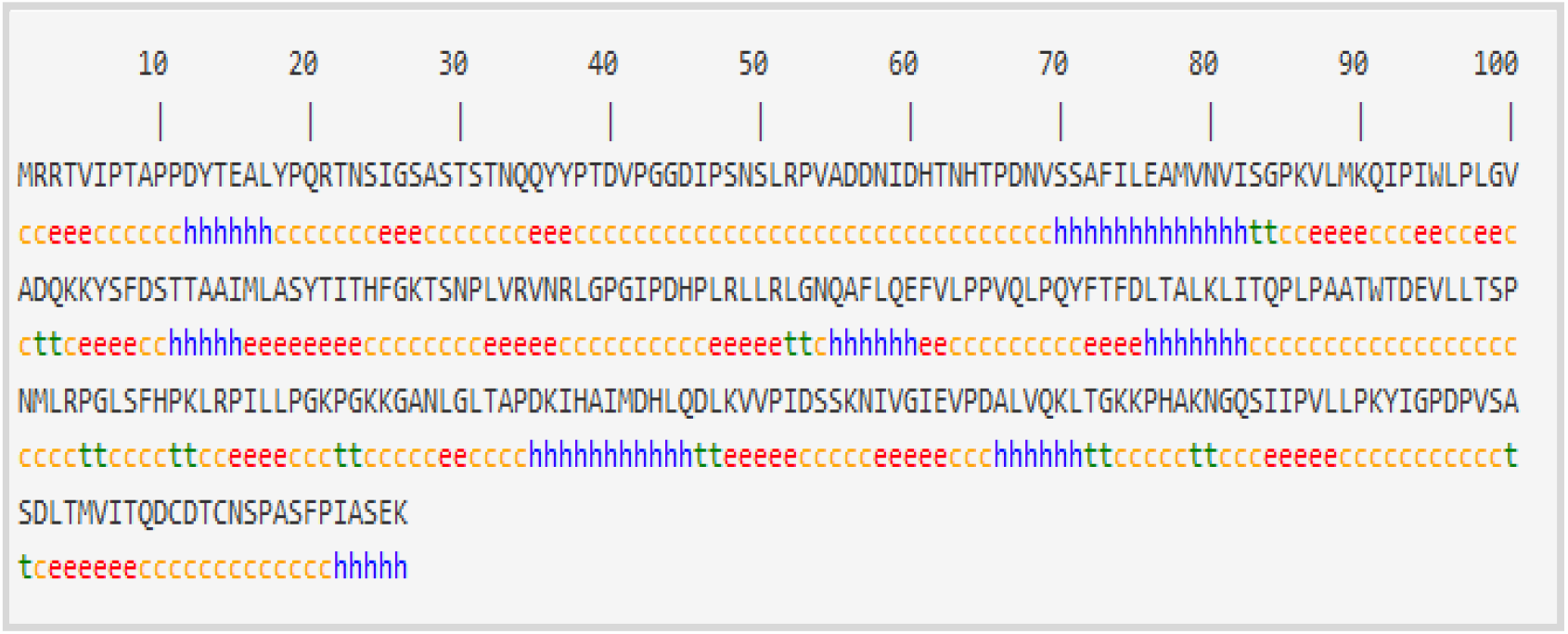
SOPMA secondary structure prediction of Bombali Ebolavirus VP40

### 3.4. 3D structure prediction and evaluation

The tertiary structure of Bombali Ebolavirus VP40 was generated using SWISS-Model sever **(Figure 5, A)**., while the Ramachandran plot created by the PROCHECK server showed that about 92% of the residues of VP40 protein are located in the most favored region **(Figure 5, B)**. Additionally, the verification of the quality of the predicted 3D structure was done using QMEAN **(Figure 6)**. QMEAN analysis resulted in a Z-score of −1.20, and the total score was 0.640. These values denote a higher quality of the model, where the acceptable score ranges between 0 and 1 [38].

**Figure 5:**
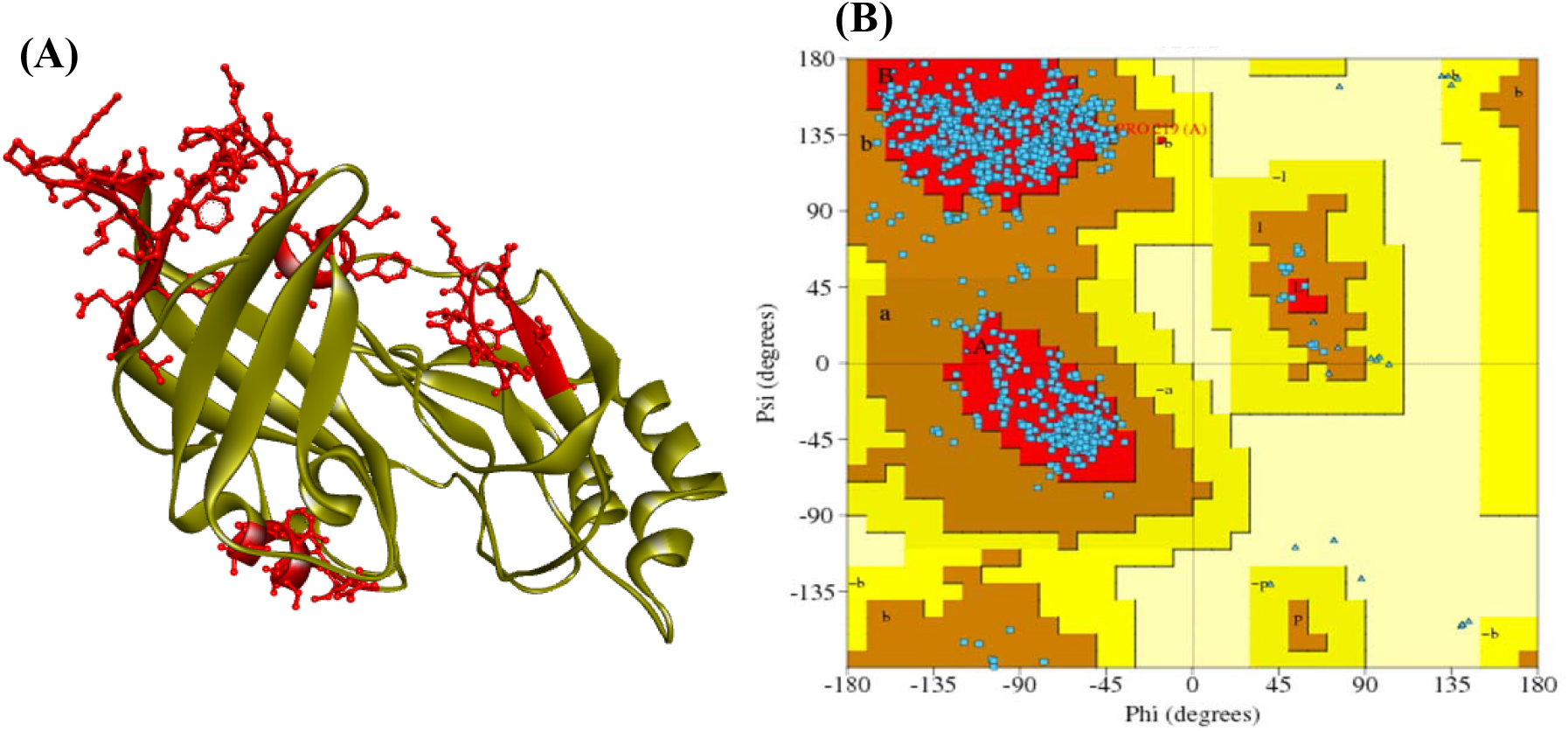
**(A)** Location of the predicted peptides (red color) within the modeled 3D structure of VP40. **(B)** Ramachandran plot analysis to validate the 3D predicted structure revealing 92% of the residues of VP40 protein are located in the most favored region.

**Figure 6:**
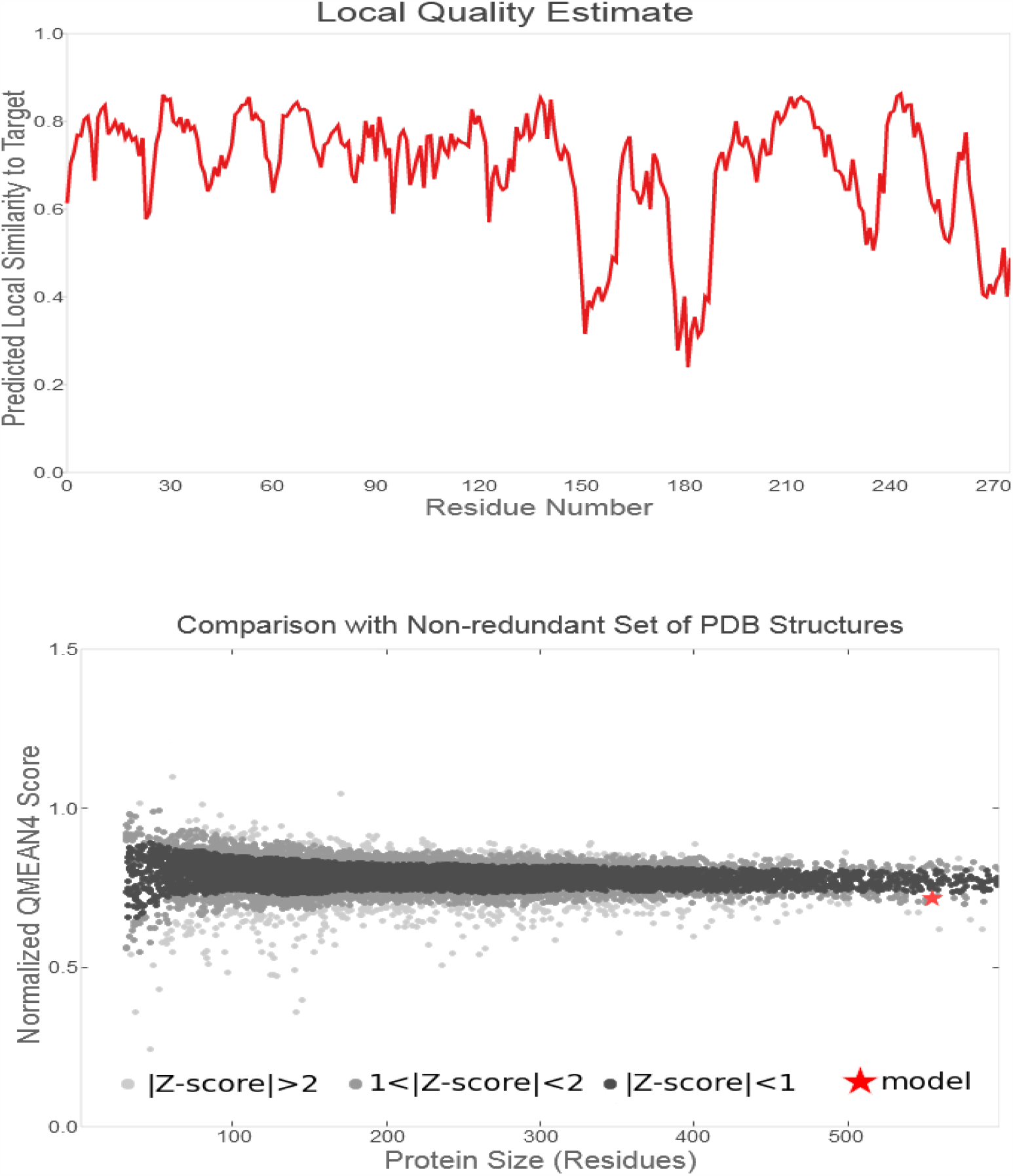
Assessment of structural superiority by QMEAN valuation.

### 3.5. T-Cell epitope prediction tools

The MHC I and MHC II binding prediction tools predicted 102 peptides and 313 peptides from the VP40 protein that could interact with different MHC I and MHC II alleles, respectively. Tables 1 and 2 summarizes the most promising peptides bound to MHC I and MHC II alleles along their allergenicity predication.

**Table 2:**
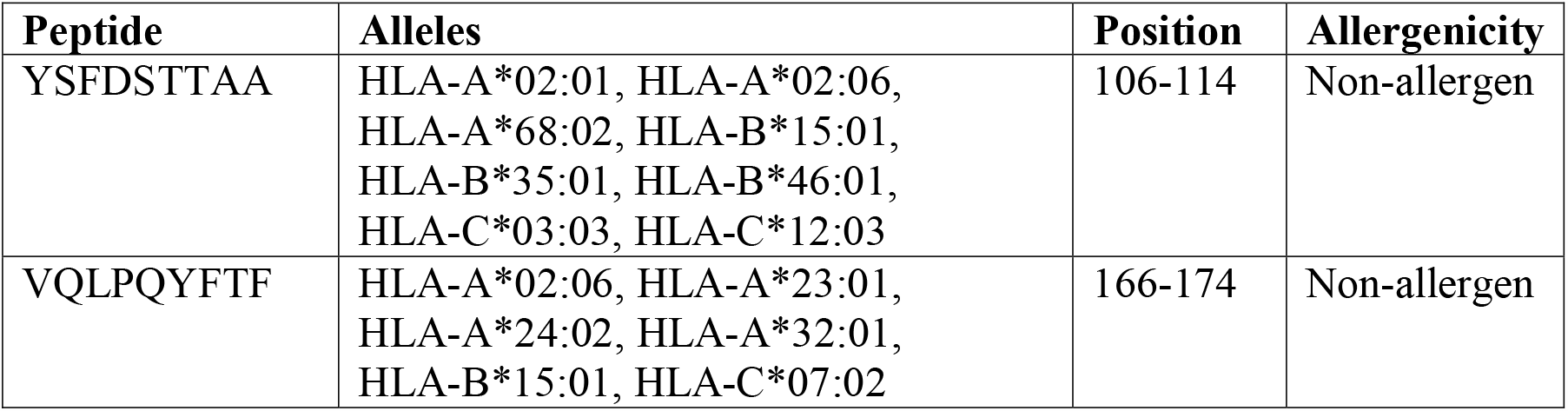
Predicted epitopes binding with MHC I alleles along with their allergenicity prediction

| Peptide | Alleles | Position | Allergenicity |
| --- | --- | --- | --- |
| YSFDSTTAA | HLA-A*02:01, HLA-A*02:06, HLA-A*68:02, HLA-B*15:01, HLA-B*35:01, HLA-B*46:01, HLA-C*03:03, HLA-C*12:03 | 106-114 | Non-allergen |
| VQLPQYFTF | HLA-A*02:06, HLA-A*23:01, HLA-A*24:02, HLA-A*32:01, HLA-B*15:01, HLA-C*07:02 | 166-174 | Non-allergen |

### 3.6. Molecular docking

**Figure 7:**
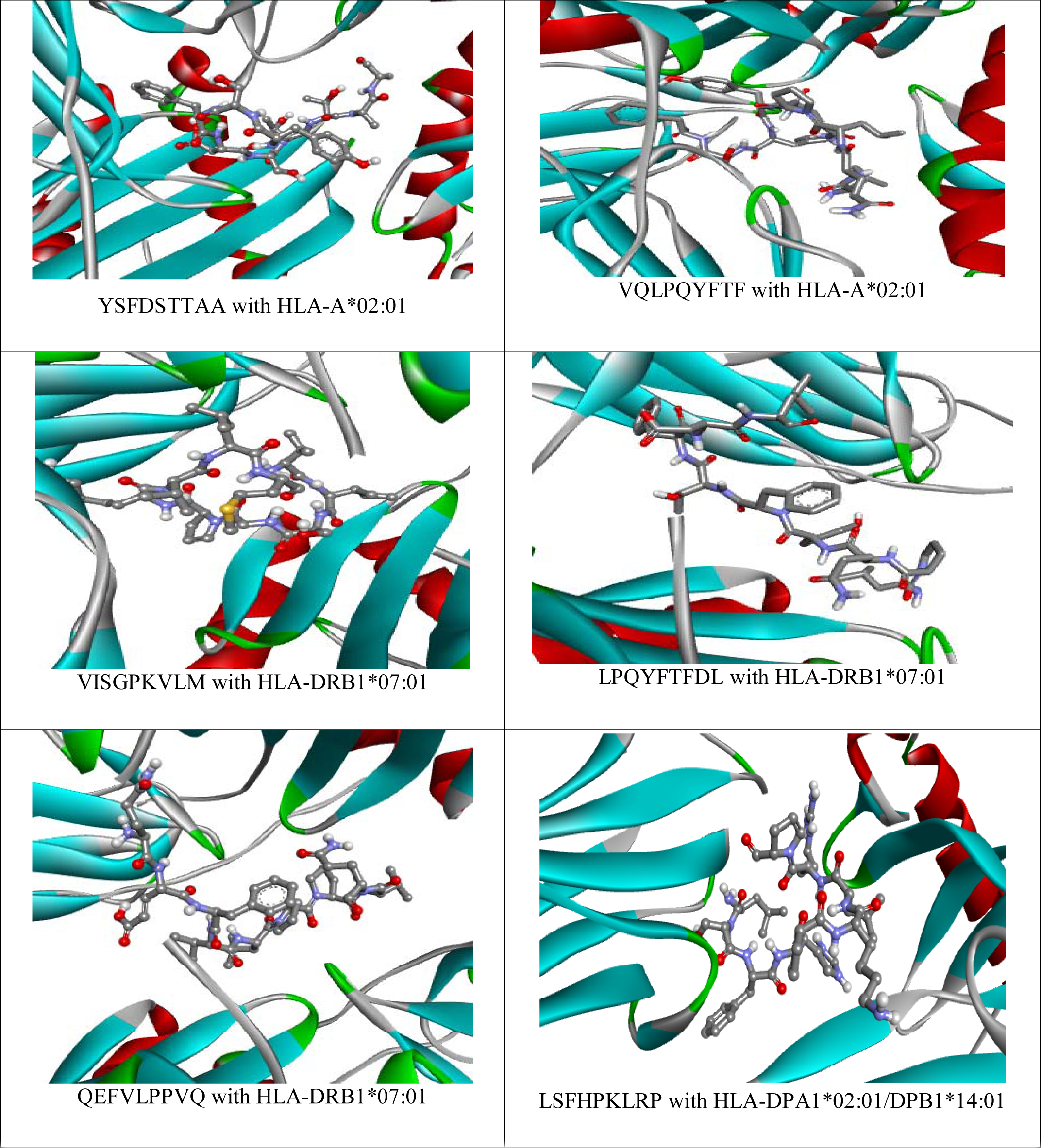
3D structures for the most promising peptides bound to MHC I and MHC II alleles.

## 4. Discussion

Ebola is a very smart RNA virus which by somehow convince the cell to engulf it, then the cell take it inside a lysosome [39]. However, Ebola needs to get out of it for two reasons, the first one is that the lysosome is filled with enzymes that can eradicates it, and all what is needed for its coping is found in the cytoplasm. The way the virus escapes is by attaching its membrane to lysosome’s membrane by receiving signals from the infected cell by identifying the right place (NPCI protein) and time to carry out this fusion reaction and release its genetic material to the cytoplasm and then initiate the replication [40, 41]. So to outsmarting ebolavirus, there are three ways to eradicates it, the first one is to go after the infected cellular machinery evolve to exploit, this is done by drug that prevent the ebolavirus from binding to NPCI; which the virus eventually get destroyed inside the lysosome because it cannot get out to replicates.

The second approach is to go after the virus itself. The glycoprotein is a highly flexible structure, thus, engineered antibody can be designed to attach to glycoprotein which will prevent the virus from binding to the host cell and eventually kill it [42-45].

The third approach is to design a vaccine which has been followed in our present study applying an immunoinformatics approach. In this study we used an immunoinformatics approach for vaccine design; we achieved protein structure-based approach for the development of multi-epitope-based peptide vaccine against Bombali Ebolavirus using matrix protein VP40 as a target.

Classifying of epitopes only based on experimental method is consuming a lot of time and resources. Recent advances of immunoinformatics methods has made it easier to identify potential epitopes, accordingly, decreasing the number of *in vivo* and *vitro* experiments for epitopes validation [18]. Peptide vaccines are more superior to traditional or single-epitope-based vaccines due to these factors: it contains multiple HLA epitopes than can be detected by many T-cell receptors; it contains multiple HLA epitopes than can be detected by many T-cell receptors; multiple epitopes provide a long-term memory, so increasing the virus range; additionally, designed vaccine can be associated with adjuvant thereby boosting its immunogenicity [18]. This study was devoted to pinpoint the 100% conserved regions which are then selected to predict the highly immunogenic epitopes for T-cells, the key molecules of cell mediated and humoral immunity as vaccine candidates for the deadly Bombali virus infection using matrix protein VP40 as a target. It’s well known that peptides can be recombinant or construct pf epitopes which are targeted against surface or intracellular proteins (such as our target) [46]. Some studies determine the ability of VP40 of Ebolavirus alone to evoke strong immune response against Ebolavirus [47, 48]. Nevertheless, there is no epitope-based peptide vaccine prediction has been done for Bombali Ebolavirus so far.

Indeed, Zaire Ebola is the strain which is responsible for most of the outbreaks with the highest case–fatality rates of all the ebolaviruses. But we are looking for the most antigenic Ebola proteins not the strain that responsible for most of outbreaks. The idea of reversed vaccinology is to replace a whole genome with small antigenic proteins [49-51], which have the ability to strongly evoke the immune system. So the antigenicity of VP40 and GP proteins from the different Ebola virus strains were analyzed by VaxiJen server **(Table 1)**. Bombali ebolavirus VP40 was found to be the most antigenic (highest score = 0.5854), therefore it was used as our vaccine target. After performing multiple alignments of all the sequences, five conserved regions formed the basis for the further analysis as they had been predicted antigenic by the VaxiJen server (Table 1).

Phylogenetic analysis reveals the evolutionary relationship between the different strains of Ebola virus and trace patterns of common ancestry between their lineages [21]. **(Figure 2)** represents maximum likelihood phylogenetic tree which constructed based on whole genome sequence alignment of the six Ebola virus strains. The taxa represent Bundibugyo Ebola (NC_014373), Tai Forest Ebola (NC_014372), Zaire Ebola (NC_002549), Bombali Ebola (NC_039345), Sudan Ebola (NC_006432), and Reston Ebola (NC_004161). Bundibugyo Ebola, Tai Forest Ebola founds on the same clade [52], also Sudan Ebola, Reston Ebola on the same clade [53] and both Zaire and Bombali Ebola are in individual clades. Zaire strain is responsible for most of the outbreaks it’s an ortholog with Bombali strain. Which supported by the high bootstrapping values on each node. As shown in **(Table 1)** the most antigenic protein among all the matrix protein of the Ebola virus, was from Bombali strains, that’s why it’s used as a target in our study.

The major histocompatibility complex (MHC) is a highly polymorphic group of cell-surface receptors found on all the cells of the body. It is well known that a strong and effective immune response depends on the recognition of epitopes by HLA molecules with high affinity. Hence, a peptide is classified by its highest number of bound HLA alleles (termed as promiscuous) has the best potential to revoke a strong immune response [54, 55].

Among the analyzed antigenic sequences, two epitopes were predicted as most promising taking into consideration their conservancy and binding affinity to the highest numbers of MHC I alleles **(Table 2)**. On the other hand, 5 conserved epitopes were predicted to interact with several MHC II alleles named HLA-D, and Q **(Table 3)**, specifying that extra attention required to be devoted to this region.

**Table 3:** Predicted epitopes binding with MHC II alleles along with their allergenicity prediction

| Peptide | Alleles | Allergenicity |
| --- | --- | --- |
| LPQYFTFDL | HLA-DPA1*01:03/DPB1*04:01, HLA-DPA1*01:03/DPB1*04:02, HLA-DPA1*01:03/DPB1*06:01, HLA-DPA1*02:01/DPB1*01:01, HLA-DQA1*01:01/DQB1*05:01, HLA-DQA1*03:03/DQB1*04:02, HLA-DQA1*01:02/DQB1*05:01, HLA-DQA1*04:01/DQB1*04:02, HLA-DRB1*04:02, HLA-DRB1*04:03, HLA-DRB1*04:04, HLA-DRB1*04:05, HLA-DRB1*07:01, HLA-DRB1*08:01, HLA-DRB1*08:02, HLA-DRB1*09:01 | Non-allergen |
| LSFHPKLRP | HLA-DPA1*02:01/DPB1*05:01, HLA-DPA1*02:01/DPB1*14:01, HLA-DQA1*01:02/DQB1*05:02, HLA-DQA1*01:02/DQB1*06:02, HLA-DQA1*02:01/DQB1*02:02, HLA-DQA1*02:01/DQB1*03:03, HLA-DRB1*01:01, HLA-DRB1*03:01, HLA-DRB1*01:03, HLA-DRB1*04:03, HLA-DRB1*04:04, HLA-DRB1*08:01, HLA-DRB1*09:01 | Non-allergen |
| QEFVLPPVQ | HLA-DPA1*01:03/DPB1*04:02, HLA-DPA1*01:03/DPB1*06:01, HLA-DPA1*02:01/DPB1*01:01, HLA-DPA1*02:01/DPB1*05:01, HLA-DQA1*01:02/DQB1*05:01, HLA-DQA1*01:03/DQB1*06:03, HLA-DQA1*04:01/DQB1*04:02, HLA-DQA1*05:01/DQB1*03:02, HLA-DQA1*06:01/DQB1*04:02 | Non-allergen |
| VISGPKVLM | HLA-DPA1*02:01/DPB1*05:01, HLA-DPA1*02:01/DPB1*14:01, HLA-DPA1*03:01/DPB1*04:02, HLA-DQA1*02:01/DQB1*04:02, HLA-DQA1*05:01/DQB1*02:01, HLA-DQA1*05:01/DQB1*03:01, HLA-DQA1*05:01/DQB1*03:03, HLA-DQA1*05:01/DQB1*04:02, HLA-DRB1*01:03, HLA-DRB1*03:01, HLA-DRB1*04:01, HLA-DRB1*04:03, HLA-DRB1*04:04, HLA-DRB1*04:05, HLA-DRB1*07:01, HLA-DRB1*08:01, HLA-DQA1*03:03/DQB1*04:02 | Non-allergen |
| YSFDSTTAA | HLA-DQA1*01:04/DQB1*05:03, HLA-DQA1*02:01/DQB1*04:02, HLA-DQA1*03:01/DQB1*03:02, HLA-DQA1*05:01/DQB1*03:03, HLA-DQA1*05:01/DQB1*04:02, HLA-DRB1*01:01, HLA-DRB1*01:03, HLA-DRB1*03:01, HLA-DRB1*04:01, HLA-DRB1*04:02, HLA-DRB1*04:03, HLA-DRB1*04:04, HLA-DRB1*08:02 | Non-allergen |

Besides the binding with the MHC molecules, the predicted peptides must be non-toxic and non-allergen, hence, their safety was predicted using AllergenFP v.1.0 [31] and AllerTop v 2.0 servers [32]. The result showed that all the selected peptides are non-allergen and non-toxin **(Tables 2, 3)**.

Population coverage analysis was also predicted for the total peptides. Obtained results showed that the proposed peptides binding to MHC class I have a 94.78% projected population coverage in the world, and 98.40% in Africa, while the population coverage results for the total number of peptides binding to MHC II alleles showed a 68.40% estimated population coverage in the world, and 85.51% in Africa.

For analyzing the stability of the vaccine, the reference sequence of Bombali Ebolavirus VP40 was used for the prediction of secondary and tertiary structure prediction by PSIPRED, and RaptorX servers respectively. A valuable insight was done from secondary structure prediction by PSIPRED. The plenty of alpha-helix and coiled region indicates high conserved regions and greater stability of VP40 protein **(Figure 3)**. To validate PSIPRED’s result, another secondary structure online tool (SOPMA) was used for this purpose **(Figure 4)**.

The tertiary structure of Bombali Ebolavirus VP40 was generated using RaptorX sever **(Figure 5, A)**, While the Ramachandran plot created by the PROCHECK server showed that about 92% of the residues of VP40 protein are located in the most favored region **(Figure 5, B)**. It is worth to mentioned that the protein model having >90% of the residues in the core and allowed regions can be designated as a high-quality model [56]. while the verification of the quality of the predicted 3D structure was done using QMEAN. QMEAN analysis, the protein model in our interest resulted in a Z-score of − 1.20, and the total score was 0.640. This value denotes a higher quality of the model, where the acceptable score ranges between 0 and 1 [38] **(Figure 6)**. On the basis of the results obtained from the mentioned structural validation tools, the 3D model showed much reliability and was considered for further study.

In order to stimulate immunological responses, the predicted peptides should interact effectively with the MHC I and MHC II molecules [57]. Therefore, molecular docking was performed to study their binding affinities selecting the alleles HLA-A*02:01, HLA-B*15:01 and HLA-DRB1 as targets due to their diversity and immunogenic association [58, 59].

H-bond formation is an important parameter to deduce the stability of the conformation through the simulation period in the protein ligand complex [60]. Thus, the PoseView at the ProteinPlus web portal was used to illustrate their 2D interactions and bonding with MHC molecules. In regards to MHC I, the obtained results showed that YSFDSTTAA and VQLPQYFTF bound to the groove of HLA-A*02:01 with binding energies of −9.3 kcal/mole and −10.2 kcal/mole, respectively. Additionally, the 2D interactions viewed that YSFDSTTAA formed eight hydrogen bonds with Asp29, Asp106, Ser4, Ser58, Arg6, Arg108, Tyr64 and Glu232. While VQLPQYFTF formed five hydrogen bonds with Lys59, Tyr64, Thr214, Glu232 and His263 **(Figure 8)**. In contrast to MHC II, the most promising peptides (LPQYFTFDL, LSFHPKLRP, QEFVLPPVQ, VISGPKVLM) were bound to MHC II molecules with binding energies of −8.4kcal/mole, −7.3 kcal/mole, −7.8kcal/mole and −6.6 kcal/mole, respectively. They formed a number of hydrogen bonds (3 to 6 hydrogen bonds) with different amino acids **(Figure 9)**.

**Figure 8:**
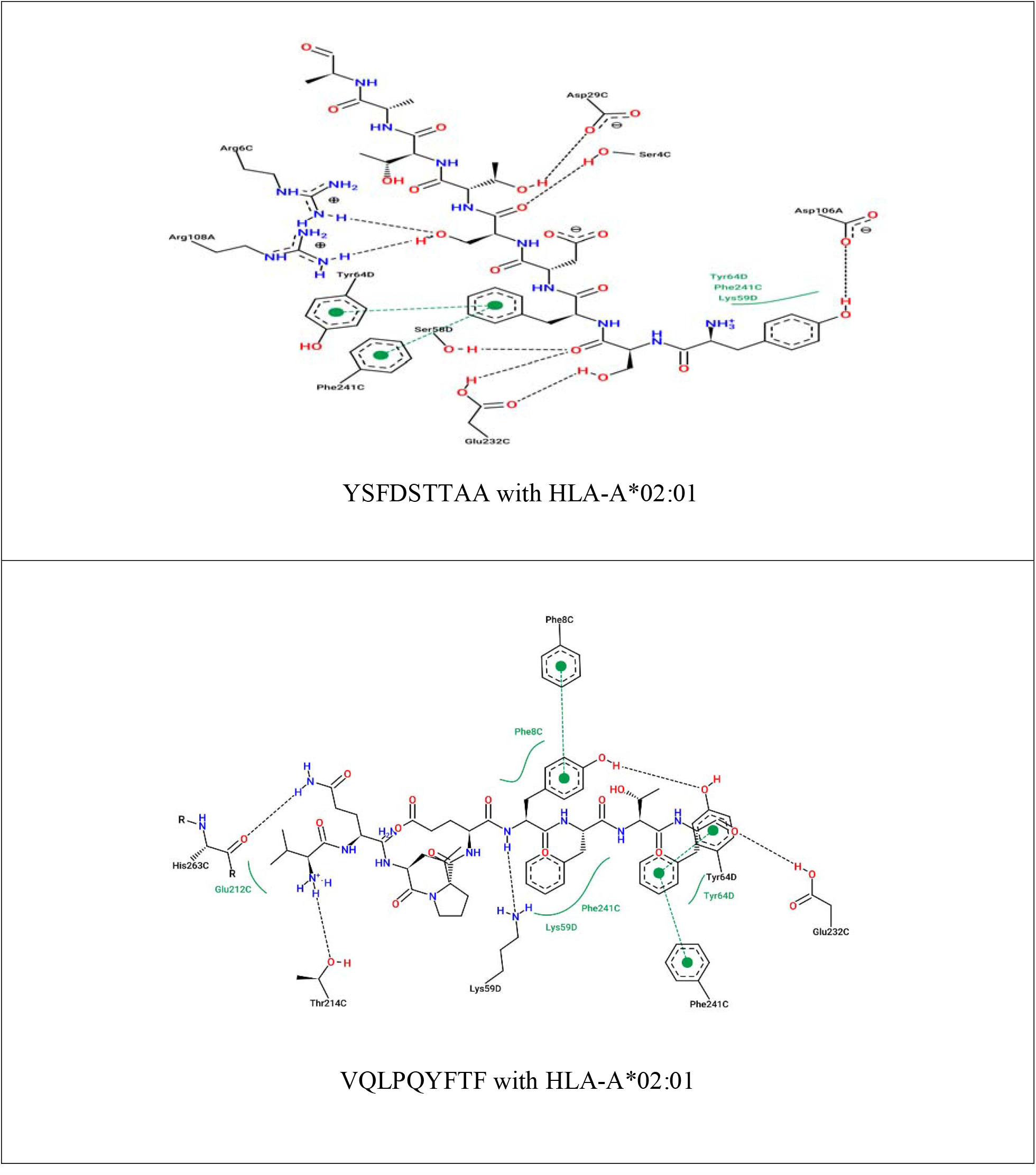
2D structure for the most promising peptides bound to MHC I alleles.

**Figure 9:**
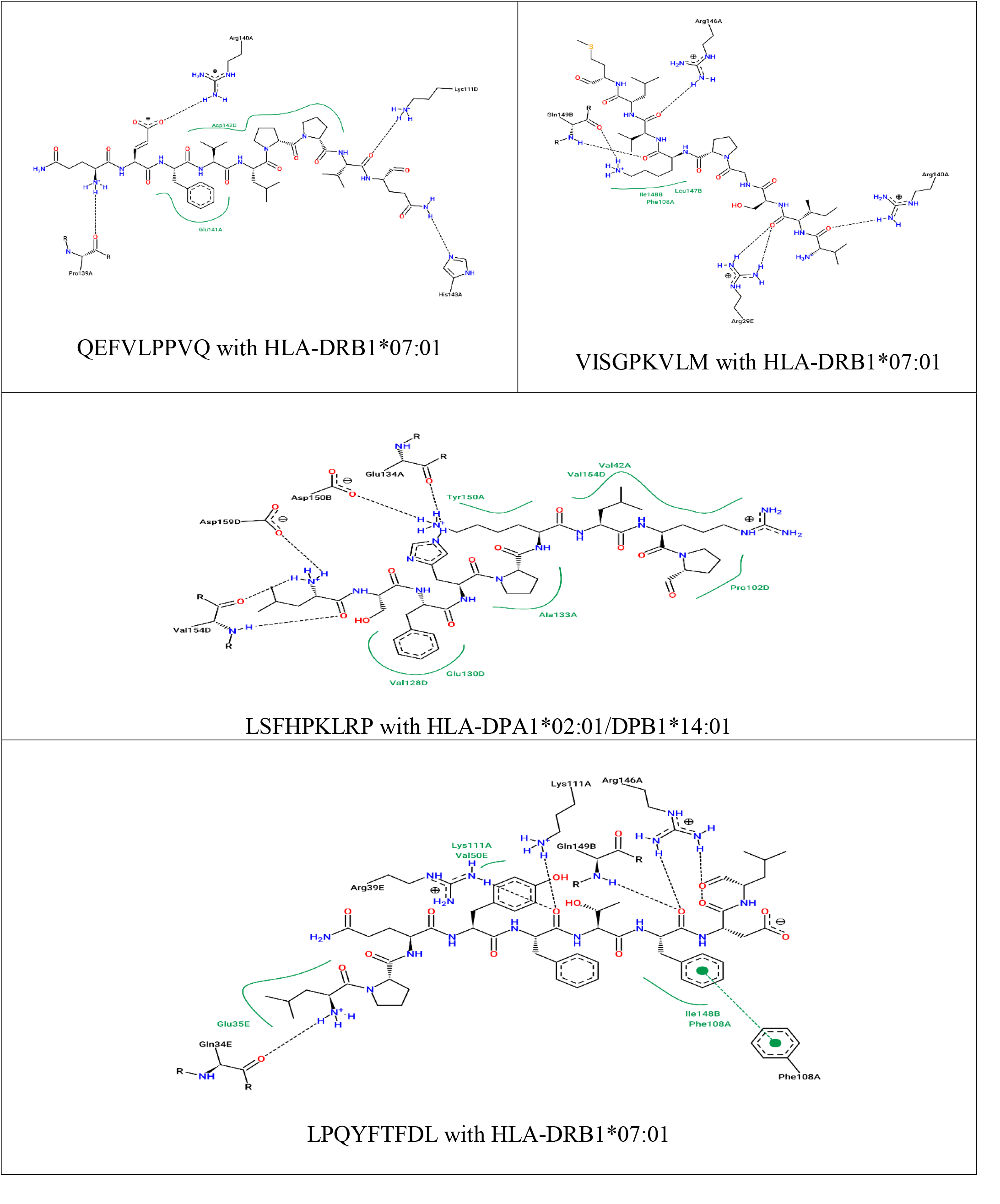
2D structures for the most promising peptides bound to MHC II alleles.

As a result of these interesting outcomes, the proposed epitopes can be used as a basis for formulating a vaccine against most of the known strains of Ebola virus, and even possibly against newly emerging strains because the basis of this study was the conservation of protein sequences in various strains.

## Conclusion

Bombali Ebolavirus is one of the most re-emerging viruses in certain regions scatter across the world. With no approved vaccine or drug against this deadly virus, the situation during an outbreak gets exacerbated. Therefore, this study devoted to serve as a platform to hasten vaccine development through design a epitope-based peptide vaccine against Bombali Ebolavirus viral Protein 40 using an immunoinformatics approaches combined with molecular docking studies. The results are originated from a systematic analysis, which suggest that, an effective vaccine candidates linking the best fit epitopes (YSFDSTTAA, VQLPQYFTF, LPQYFTFDL, QEFVLPPVQ, VISGPKVLM, and LSFHPKLRP) and the complete analysis of this proposed vaccine discloses the high tendency of the vaccine to provoke a strong immune response against Bombali Ebolavirus using matrix protein VP40 as a target. Finally, both in vivo and in vitro experiments are suggested to support these findings.

## Data Availability

All data underlying the results are available as part of the article, and no additional source data are required.

## Authors’ Contributions

MIM: conceptualization, formal analysis, methodology, validation, and writing (original draft); SWS: conceptualization, formal analysis, visualization, validation, and writing (original draft); MIA: conceptualization, formal analysis, methodology, writing (original draft); and AMM: data curation, conceptualization, project administration, supervision, and writing (review and editing). All authors have read and approved the final manuscript.

